# Improved cell-free transcription-translation reactions in microfluidic chemostats augmented with hydrogel membranes for continuous small molecule dialysis

**DOI:** 10.1101/2022.08.23.504913

**Authors:** Barbora Lavickova, Laura Grasemann, Sebastian J Maerkl

**Affiliations:** Institute of Bioengineering, School of Engineering, École Polytechnique Fédérale de Lausanne, Lausanne, Switzerland

## Abstract

Increasing protein production capacity of the PURE cell-free transcription-translation (TX-TL) system will be key to implementing complex synthetic biological circuits, and to establish a fully self-regenerating system as a basis for the development of a synthetic cell. Under steady-state conditions, the protein synthesis capacity of the PURE system is likely at least one order of magnitude too low to express sufficient quantities of all PURE protein components. This is in part due to the fact that protein synthesis can’t be sustained during the entire dilution cycle, especially at low dilution rates. We developed a microfluidic chemostat augmented with semi-permeable membranes that combines steady-state reactions and continuous dialysis as a possible solution to enhance protein synthesis at steady-state. In batch operation, the continuous dialysis of low molecular weight components via the membranes extended protein synthesis by over an order of magnitude from 2 hours to over 30 hours, leading to a seven-fold increase in protein yield. In chemostat operation, continuous dialysis enabled sustained protein synthesis during the entire dilution cycle even for low dilution rates, leading to six-fold higher protein levels at steady state. The possibility to combine and independently manipulate continuous dialysis and chemostat operation renders our dialysis chemostat a promising technological basis for complex cell-free synthetic biology applications that require enhanced protein synthesis capacity.

## Introduction

A reconstituted cell-free transcription-translation (TX-TL) system such as the PURE system is a viable chassis for constructing a synthetic cell^1^. However, one critical requirement that remains a major challenge is achieving a sufficiently high protein synthesis rate to regenerate all PURE proteins simultaneously in order to achieve sustained self-regeneration. We recently demonstrated that several proteins of the PURE system could be sustainably self-regenerated in a microfluidic-based synthetic cell-like system^2^. However, the synthesis rate and yield in the recombinant cell-free system are currently predicted to be orders of magnitude below what would be required to regenerate all of its non-ribosomal protein components, which is a limitation not only for advances in constructing a synthetic cell, but also for prototyping of complex synthetic systems.

Cell-free reactions can be classified into equilibrium and non-equilibrium reactions^1^. Standard batch experiments are classified as equilibrium reactions. All components are combined at the outset, after which they undergo reactions with no further external input, until chemical equilibrium is reached. A standard cell-free protein synthesis system in batch configuration ceases to function mainly due to the rapid depletion of critical small molecular weight components such as NTPs^1^. This, however, does not mimic biological systems, which operate far from chemical equilibrium. Therefore, continuous in vitro systems, where reagents are exchanged over time, have been developed.

These non-equilibrium reactions can be further separated into two sub-categories: batch reactions with continuous dialysis, and reactions in chemostat reactors. During batch reactions with dialysis, replenishment is achieved by passive, diffusion driven exchange based on dialysis, which allows the exchange of small molecules with the environment, while retaining the TX-TL machinery within a defined reaction compartment^3^. This approach permits protein synthesis to last for up to several days, leading to a total protein synthesis yield in the range of mg/mL^4^. Batch reactions with continuous dialysis have previously been realized using standard tube-based dialysis^4^, a micro-well plate with integrated dialysis membranes^5,6^ or a passive PDMS microreactor^7^. In comparison, reactions in chemostat reactors overcome equilibrium by diluting all reaction components with fresh components; this allows for replenishment of both small molecules and the enzymatic machinery, allowing an efficient protein turnover rate and thus implementation of complex genetic networks^8–10^. Therefore, combining both approaches by replenishing small molecules by dialysis and replenishing the enzymatic machinery by dilution is likely crucial to achieve adequate synthesis rates required for self-replicating systems.

In this work, we designed a microfluidic device with 8 independent microfluidic chemostat reactors with integrated PEG-DA dialysis membranes. Our device enables the implementation of steady-state reactions in combination with small molecule feeding by continuous dialysis. We describe a simple silanization protocol that does not require either the use of oxygen plasma or organic solvents, and a technique for simple PEG-DA hydrogel membrane patterning using pneumatic valves in an oxygen-free environment. Additionally, we describe how different membrane permeabilities can be generated to achieve an ideal molecular weight cut-off for a continuous dialysis TX-TL reaction. Furthermore, we report a simple way to avoid water evaporation from PDMS at low environmental humidity by adding a PDMS hydration layer. We show that implementing continuous dialysis protein synthesis in our steady-state chemostat reactor with semi-permeable membranes extended protein synthesis to at least 30 hours, leading to a sevenfold increase in the synthesis yield compared to batch reactions. Moreover, by combining the chemostat TX-TL reaction with continuous dialysis, we achieved a six-fold increase in steady-state protein synthesis levels while maintaining active protein turnover.

## Results

### Formulation, generation, and characterization of PEG-DA hydrogel membranes

To achieve continuous dialysis of small molecules, several methods and designs have been proposed ^5,6,7^, with most of them suffering from the drawback that they are difficult to implement and manufacture. We adapted a method to manufacture PEG-DA semi-permeable membranes inside of a microfluidic device utilized for protein crystallization^12^. PEG-DA is ideal for this purpose due to several advantages including biocompatibility^13^, tunable pore sizes^11,14–16^, the possibility to be integrated within PDMS^12^, the ability to be introduced as the unpolymerized prepolymer, and to be quickly cured by UV. The microfluidic reactor design used fluidically hard-coded dilution fractions as described previously^17^, and was augmented with a feeding channel for the continuous dialysis of small molecules, and an anti-evaporation layer on top of the device (Supplementary Figure 1a-f). The feeding channel is linked to the main reactor by four channels, which contain the PEG-DA hydrogel membranes. These PEG-DA membranes form a semi-permeable barrier between the reactor and feeding chamber to allow the continuous supply of small molecular weight components while retaining high molecular weight components such as proteins or ribosomes inside the reactor (Figure 1a,b, Supplementary Figure 1a,b). These linker channels can be controlled by two valves on either end. To generate PEG-DA membranes, the valves facing the reactor ring were closed to prevent any chemical from entering the actual reaction chamber, and the linker channels were addressed via the feeding chamber. The PDMS surface was functionalized with silane as described below, and subsequently PEG-DA was flown into the channels. The valve facing the feeding chamber was actuated to trap the PEG-DA inside the linker channels, and the feeding chamber was thoroughly washed to remove all remaining PEG-DA. The PEG-DA was then cured using a UV lamp generating semi-permeable membranes separating the main reactor from the feeding channel. To implement the semi-permeable hydrogel membranes for the application of long-term continuous dialysis of small molecular reagents during cell-free expressions, several crucial factors, such as strong anchoring of the membranes to the PDMS and glass surfaces, a correct molecular weight cut-off, and a low evaporation rate of liquids from the reactors had to be achieved.

**Figure 1.**
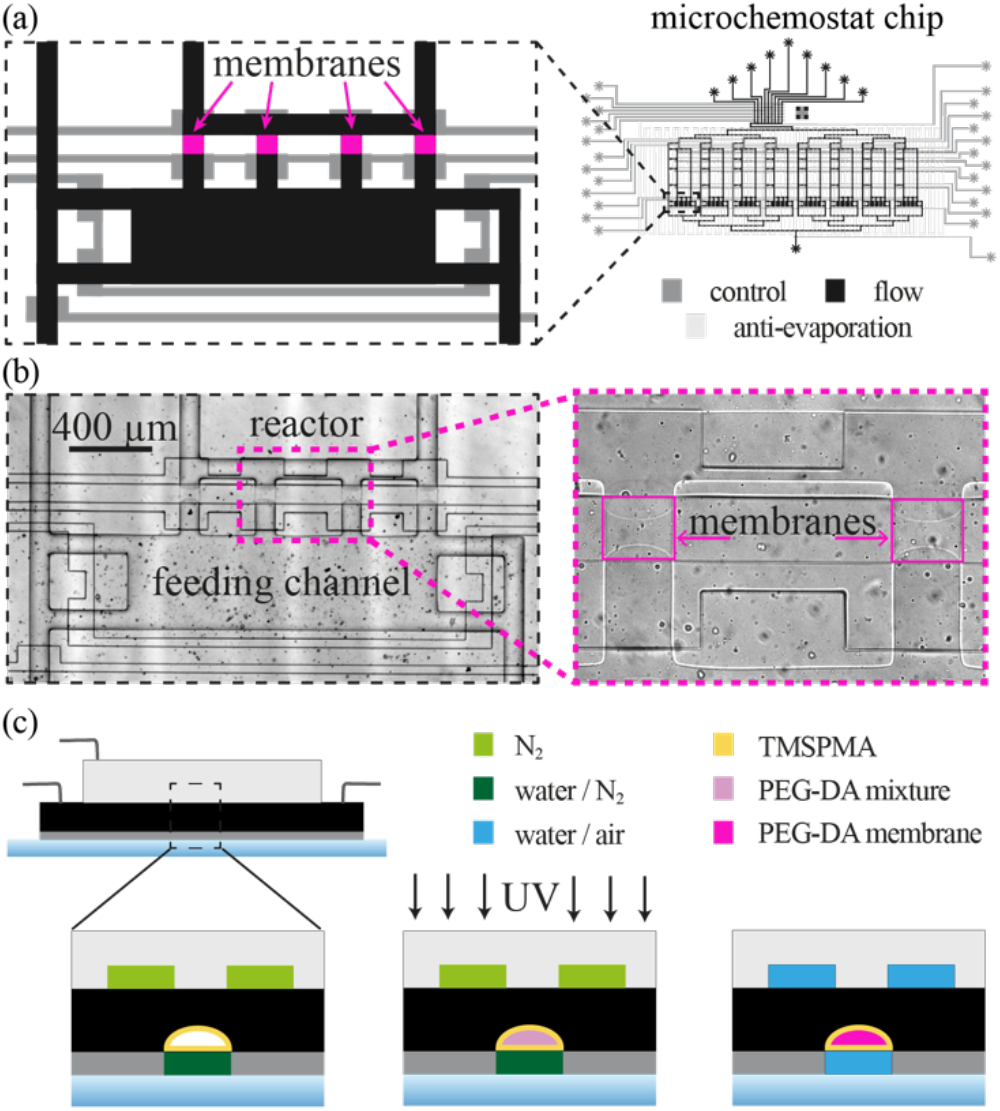
(a) Design schematic of the microfluidic device featuring eight individual chemostat reactors and details of the bottom of a single reactor showing the feeding channel separated from the main reactor by hydrogel membranes. The flow layer is shown in black, the control layer in dark gray, and the anti-evaporation layer in light gray. Design and functional details are provided in Supplementary Fig. 1. (b) Bright-field image of the hydrogel membranes between the reactor and feeding channel. (c) Schematic protocol for preparing hydrogel membranes. First the PDMS channels were functionalized with TMSPMA, subsequently PEG-DA prepolymer was flown and polymerized inside of the membrane channels, and the formed membranes were washed overnight with water before conducting on-chip cell-free TX-TL reactions.

In our device, pressure on the membranes is imposed over extended periods due to the osmotic pressure generated by the differences in composition of the feeding solution and the PURE TX-TL solution, as well as during the pressure-driven loading phase, and as a result of volume displacement arising from the pneumatic valve actuation. Therefore, strong anchoring of the membranes to PDMS is crucial. To anchor the membranes, the addition of vinyl groups to the PDMS surface by silanization with silane-containing acrylate or methacrylate^18^ was essential. Here we report a simple and fast (< 1h) silanization protocol based on hydrochloric acid/hydrogen peroxide treatment^19,20^ and TMSPMA hydrolyzed in acidic conditions^21^, without the use of organic solvents, which can support strong PEG-DA anchoring and is compatible with PDMS valves (Figure 1c). Polymerization of PEG-DA membranes has to be performed in an oxygen-free environment, as the presence of oxygen leads to a thin oxidized layer that is not able to cross-link with the PDMS^12^. To deplete oxygen, the hydrogel-forming valves controlling the connection of the feeding channel with the ring, and the anti-evaporation layer were pressurized with nitrogen instead of air (Figure 1c). The resulting channel functionalization and formation of hydrogel membranes was highly reproducible, and the membranes were stable against rupture and could withstand the imposed trans-membrane pressure for more than two days.

To achieve continuous cell-free expression, membranes of the correct molecular weight cut-off have to be implemented on the device. The ideal molecular weight cut-off level for cell-free continuous dialysis reactions is around 10 kDa^12^, which allows small molecules to diffuse between feeding and reaction chamber, while retaining protein components inside the reaction chamber. One of the main advantages of utilizing hydrogel membranes as dialysis membranes is their tunable pore size. In particular PEG-DA hydrogels have been shown to be tunable from nm pores^22^ to μm^14–16^ pores, allowing molecular weight cut offs from small molecules to large proteins. The porosity depends mostly on the molecular size of the PEG-DA polymer^14^ and porogen^15,22^ used during the polymerization reaction. To achieve the desired permeability of our membrane and low hydrogel swelling^23,24^, we decided to use PEG-DA M_n_ 700 g/mol (average M_n_) with photo-initiator 2-hydroxy-2-methylpropio-phenone^12^. The main advantage of using M_n_ 700 g/mol PEG-DA prepolymer is its solubility in water, in contrast to M_n_ 250 g/mol PEG-DA prepolymer for example, which is insoluble in water. This significantly simplifies prepolymer manipulation within the microfluidic chip, as any remaining unpolymerized prepolymer can easily be washed away. Moreover, water simultaneously acts as a porogen during hydrogel polymerization, therefore the hydrogel permeability can be tuned by modifying the PEG-DA/water ratio.

We initially prepared two aqueous formulations of different PEG-DA/water ratios (33 % and 42 % w/w PEG-DA/water), and examined the diffusion of different molecular weight solutes through the formed membranes from the feeding channels to the chemostat reactor (Figure 2a, b). While the small molecular weight methylene blue dye (Figure 2c) diffused rapidly into the reactor for both tested PEG-DA formulations, 10 kDa (Figure 2d) and 40 kDa (Figure 2e) fluorescently labeled dextrans were mainly retained in the feeding channel, and diffused into the reactor at a much slower rate. In particular for the membrane formulation of 42 % w/w PEG-DA/water, there was no observable diffusion into the reactor ring for the 10 kDa or the 40 kDa dextran in the time frame of the experiment. This renders both formulations ideal candidates to be utilized for membranes for continuous dialysis protein synthesis, and we decided to implement the 33 % w/w PEG-DA/water formulation for subsequent experiments.

**Figure 2.**
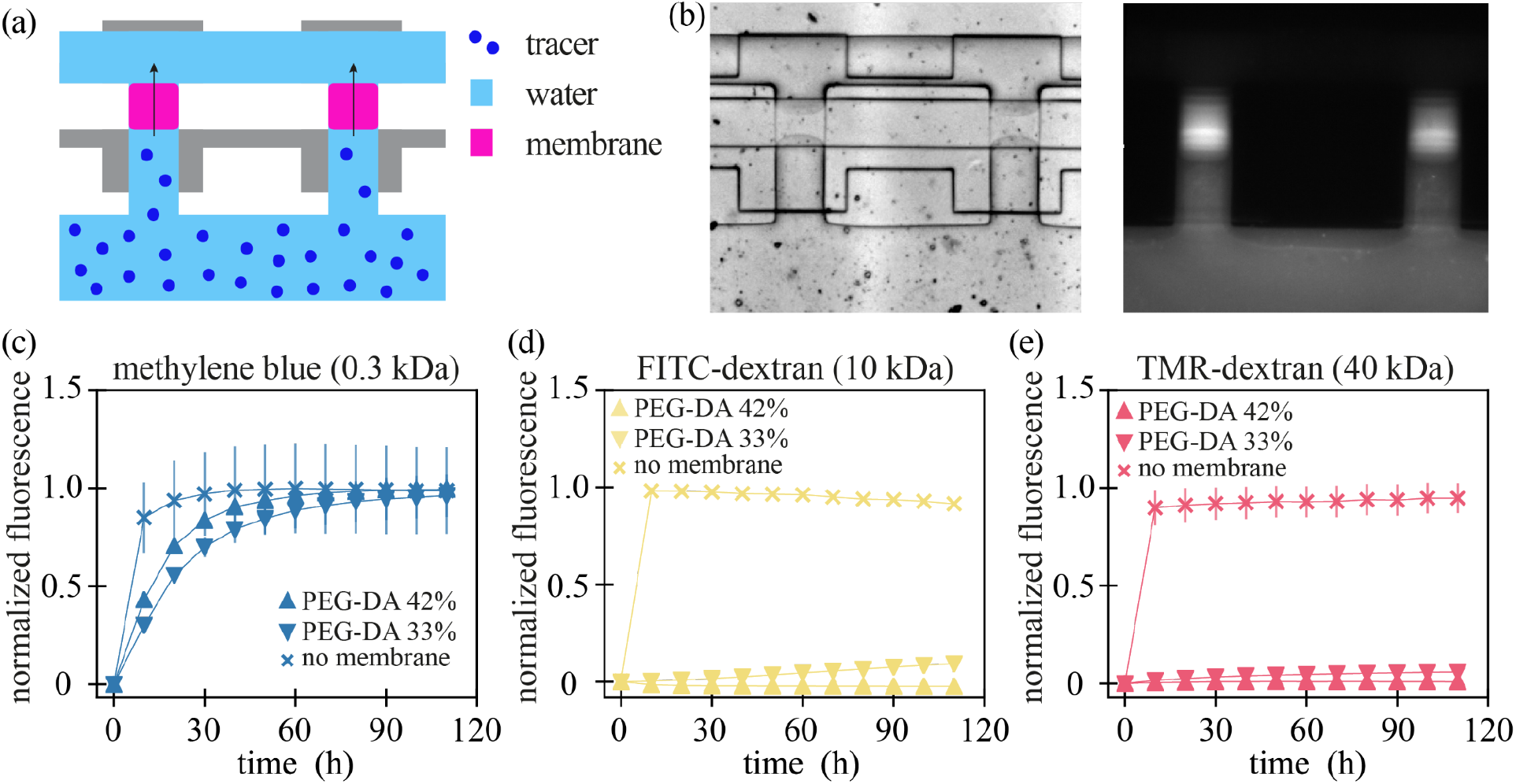
Solute diffusion through the membrane: (a) Schematic of a hydrogel permeability experiment. The feeding chamber (bottom) and the reaction ring (top) are separated by semi-permeable membranes (pink) which allow small molecules to diffuse freely, while retaining molecules above the respective molecular weight cut-off. (b) Bright field and fluorescent images of the semi-permeable membrane and the feeding channel loaded with methylene blue dye. Diffusion of solutes of different molecular weight from the enclosed feeding channel to the reactor chamber through the membrane over time: (c) methylene blue, (d) 10 kDa FITC-dextran, (e) 40 kDa TMR-dextran. The fluorescence of methylene blue was normalized to the maximum level attained in each experiment. For 10 kDa FITC-dextran and 40 kDa TMR-dextran, the fluorescence was normalized to the maximum level attained in reactors without membrane.

### Batch cell-free expression augmented with continuous dialysis

We first tested if our microfluidic device with integrated membranes could be used for batch cell-free protein synthesis and whether the dialysis membranes have a noticeable effect on protein synthesis. To be able to sustain long-term protein synthesis we implemented an anti-evaporation layer to prevent evaporation of water through the PDMS. In a PDMS device during a batch reaction (environmental humidity and 34 °C), evaporation can be observed after 2 hours, if no evaporation prevention is applied (Supplementary Figure 1g). The anti-evaporation layer features a dead-end serpentine channel covering the reactors, and is bonded on top of the chip using oxygen plasma (Supplementary Figure 1 c-f). During the experiment, the channel is connected to a pressurized, water-filled tube to sustain a constant water supply in the anti-evaporation layer. With the anti-evaporation layer, no discernible evaporation was observed after 16 hours (Supplementary Figure 1g). Compared to previously reported anti-evaporation methods^25–28^, our design is easy to fabricate, assemble, and can be applied to most standard double-layer microfluidic designs without the need of additional hardware.

For cell-free expression experiments, the chemostat reactors were filled with the PURExpress components in a ratio of 2:2:1 for solution A (2.5x, energy solution), solution B (2.5x, protein/ribosome), and DNA solution (5x), respectively. The feeding channel was loaded with a 1.5x feeding solution (1.5x solution A and PURE buffer (see methods)), either once in the beginning of the experiment for the batch with static dialysis reaction, or every 10 min for the continuous dialysis experiments (Figure 3a). The batch reaction was run as previously described^2^ using a traditional chemostat device without a feeding chamber, but augmented with the anti-evaporation layer. The experimental procedures and parameters of the batch, batch with static dialysis, and batch with continuous dialysis experiments are detailed in Supplementary Table 1.

**Figure 3.**
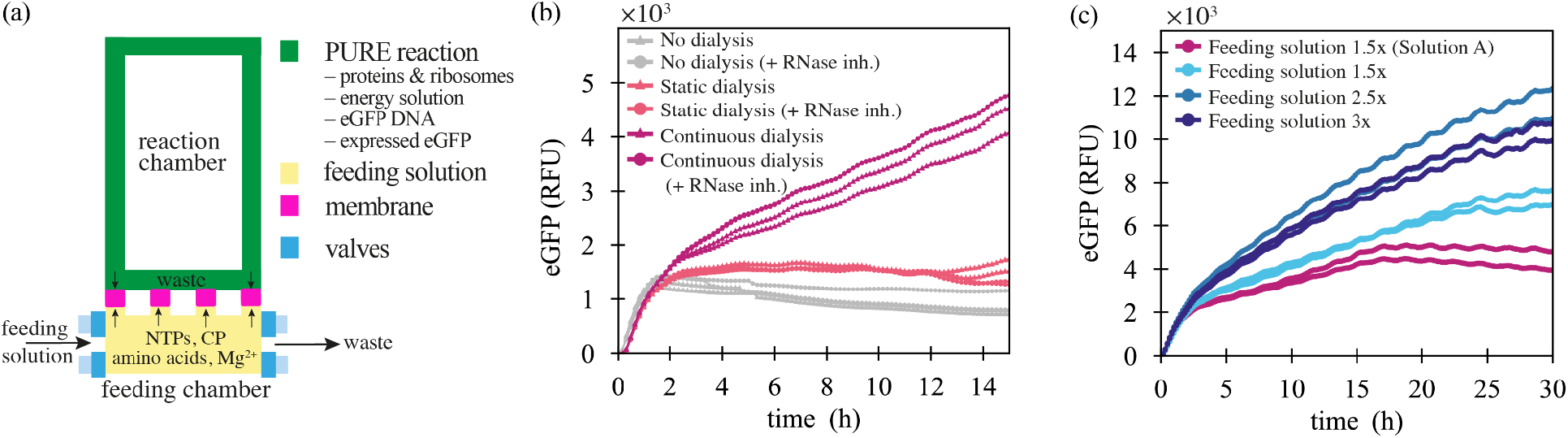
Batch cell-free expression augmented with dialysis: (a) Schematic of a continuous dialysis reaction of a single chemostat reactor augmented with a hydrogel membrane. (b) Time course of in vitro eGFP expression in batch, batch with static dialysis, and batch with continuous dialysis reactions. A concentration of 1.5x of the feeding solution based on solution A was used for batch with static dialysis and batch with continuous dialysis reactions. Each curve represents a technical replicate. (c) Time course of in vitro eGFP expression levels in batch with continuous dialysis reactions for different feeding solution concentrations based on the commercial solution A (PURExpress) or home-made energy solution. Each curve represents a technical replicate.

There was no significant difference between the batch reaction and batch with static dialysis reaction, as the feeding channel was filled only once at the beginning of the reaction, therefore supplying only a limited amount of additional energy components. For both reactions, protein synthesis ceased after around 2 hours. However, continuous eGFP synthesis was observed for the batch reaction with continuous dialysis, when the feeding solution was replenished every 10 min, which resulted in a three-fold increase in the reaction yield after 15 hours compared to the batch reaction with static dialysis (Figure 3b). In addition, we explored the importance of an RNAse inhibitor to prevent the degradation of RNA based components, but did not observe any difference between the reactions with and without RNAse inhibitors for any conditions tested (Figure 3b).

We tested different concentrations of the feeding solution to further increase protein synthesis (Figure 3c, Supplementary Figure 2). We prepared different concentrations of feeding solutions, with energy solutions prepared based on previously published protocols^29^, and compared them to the feeding solution based on the commercially available solution A of the PURExpress system. We achieved a similar expression level comparing our homemade 1.5x solution to the 1.5x concentrated PURExpress solution A for up to 15 hours. However, in contrast to the homemade solution, which exhibited steady protein expression during the entire experiment of 30 h, we observed a cessation of eGFP expression after 15 hours when using the commercial solution. This effect is potentially due to the use of the more stable TCEP as a reducing agent in the homemade solution instead of DTT that is used in the commercial solution. Increasing feeding solution concentration to 2.5x led to an increase in synthesis rates and overall higher yield by around two-fold compared to the 1.5x feeding solution concentration. After 30 hours, we achieved a seven-fold increase of protein yield without observing any cessation of protein synthesis. Further increasing the concentration (3x) did not increase synthesis rate or yield compared to the 2.5x feeding solution.

### Steady-state cell-free reaction augmented with continuous dialysis

A system that is designed to implement complex biological networks needs to be able to exchange reagents and operate at steady-state. We have previously shown that a microfluidic chemostat can achieve steady-state transcription and translation^2,9,10^. However, a remaining challenge is obtaining adequate protein synthesis at steady-state to implement larger and more complex biological circuits and to establish a self-replicating system for a future synthetic cell. Steady-state is achieved when the synthesis rate matches the combined dilution and degradation rate. Hence, at a given synthesis rate, faster dilutions lead to lower steady-state levels, and vice versa. In the experiments described in this work, the fraction of the ring being exchanged is held constant at 20%, and the dilution rate is determined solely by the dilution frequency. In silico models of the chemostat show that in the ideal case, where the synthesis rate remains constant throughout the time interval between dilution steps, steady-state levels are inversely proportional to dilution rates (Supplementary Figure 3a). For instance, a dilution interval of 60 min should lead to a four-fold increase in steady-state GFP level compared to a 15 min dilution interval. However, if the supply of small molecules is limiting (Supplementary Figure 3b) or inhibitory byproducts accumulate, the synthesis rate between dilution steps may not be constant and decrease or cease entirely, and hence lower dilution rates will not lead to proportionally increased steady-state protein levels.

To test whether resource supply might be a limiting factor, as we hypothesized in our previous work^2^, we investigated steady-state expression at different dilution rates. The chemostat reactors were loaded with PURExpress components in a ratio of 2:2:1 (v/v/v) for energy solution (2.5x, energy components), solution B (2.5x, proteins/ribosomes), and DNA solution. Subsequently, to investigate steady-state expression levels at different dilution rates, 20 % of the reactor volume was exchanged with fresh components using the same ratio every 15, 30, or 60 min (Supplementary table 2.). We observed similar steady-state levels for all dilution frequencies, with the lowest steady-state level for the 60 min dilution interval. This indicates that the synthesis rate averaged over one cycle is the lowest for the 60 min dilution interval. As it can be seen in Figure 4a, the synthesis rate in fact decreases during the cycle, probably due to insufficient supply of energy components. As lower dilution rates mean less components being replenished per unit time, and the 20 % volume replacement every 60 min is apparently insufficient to supply enough small molecules to sustain a constant synthesis rate during the 60 min reaction, a lower steady-state protein level is achieved.

**Figure 4.**
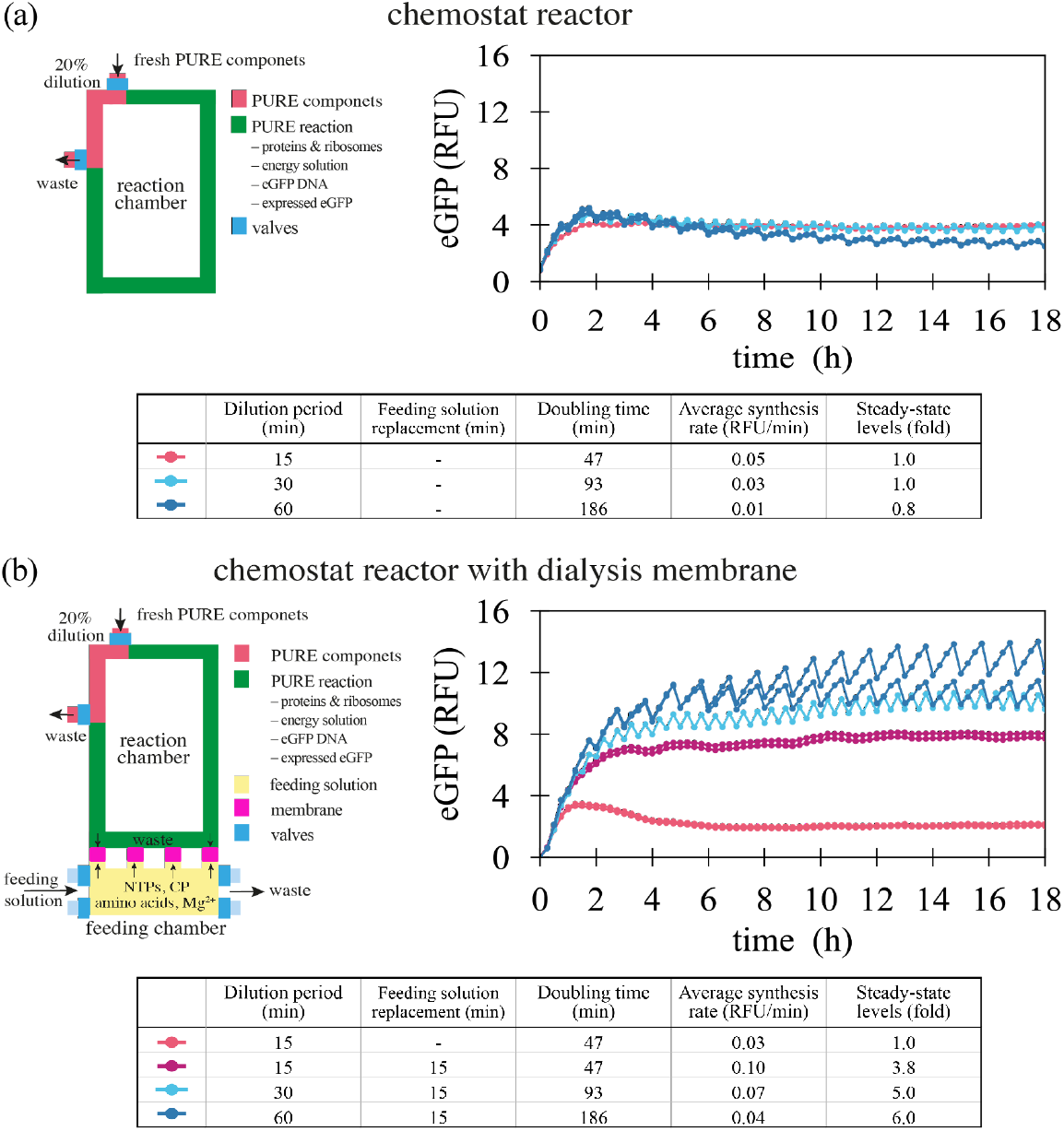
Steady-state chemostat reactions. (a) eGFP steady-state levels on a traditional chemostat reactor as previously described30 for different dilution rates. 20% of the chemostat reactor volume was diluted with energy solution, solution B (PURExpress), and DNA at different time intervals of 15, 30, and 60 min. Each curve represents a technical replicate. (b) eGFP steady-state levels for chemostat reactors augmented with hydrogel membranes for different dilution rates. 20% of the chemostat reactor volume was diluted with energy solution, solution B (PURExpress), and DNA at different time intervals of 15, 30, and 60 min. The feeding solution (3x) was replaced as specified in the table. Each curve represents a technical replicate. Doubling time t_d_ = ln(2) μ^−1^ was calculated from the dilution rate μ = −ln(C_t_/C_0_)t^−1^. The average synthesis rate for each dilution period was calculated from the steady-state levels (average during 5-18h) based on the percentage of dilution and the dilution period.

We thus combined the chemostat reaction with a continuous dialysis reaction to overcome the insufficient supply of small molecular components at lower dilution rates (Figure 4b). For continuous dialysis reactions, the chemostat reactors were filled and diluted as described above for the standard chemostat reactions. In addition, the feeding channel was loaded with a 3x feeding solution (3x energy solution and PURE buffer, see methods), which was exchanged every 15 min, if not indicated otherwise (Figure 4b). Surprisingly, we already observed a significantly enhanced eGFP synthesis rate (four-fold) when continuously exchanging the feeding solution for the 15 min dilution interval, indicating that even our highest dilution rate is already resource limited. By lowering the dilution rates, we further increased the steady-state level eGFP fluorescence level by six-fold compared to the reaction without continuous dialysis. In contrast to the traditional chemostat reactor, we did not observe any significant decrease in synthesis rate during each dilution cycle even for the longest dilution interval (60 min), indicating a sufficient supply of small molecules through the membranes to sustain protein expression. However, the average synthesis rate for the 60 min interval was lower than the average synthesis rate at an interval of 15 min, suggesting that further optimization might be possible.

## Discussion

In this work, we augmented a microfluidic reactor with hydrogel-based semi-permeable dialysis membranes to combine chemostat operation with continuous dialysis. For standard batch reactions augmenting the reaction chambers with semi-permeable membranes led to a prolongation of protein synthesis from approximately two to at least 30 h, increasing total protein yield by seven-fold. For chemostat operation we showed that combining steady-state protein synthesis with continuous dialysis led to a six-fold increase in protein levels at steady-state. By decoupling small molecule supply from TX-TL machinery replenishment and dilution, we more closely mimic the dilution and regeneration of cellular components during cell growth. In a living cell, small molecules are taken up from the outside to fuel reactions inside the cell, providing a supply of small metabolites that is decoupled from growth and thus dilution. Large biomolecules including proteins and DNA are synthesized inside the cell and diluted (or rather kept constant) by growth and degradation. Likewise, our chemostat exchanges small molecules with the feeding chamber to fuel the reactions inside, while keeping proteins and DNA inside the reactor subject to exchange by dilution. To our knowledge, this is the first device that combines chemostat operation and continuous dialysis cell-free protein expression.

The increase in the protein production capacity at steady-state is a key step towards achieving adequate protein synthesis likely required when building a synthetic cell based on the PURE cell-free expression system. We anticipate that further optimizing and tuning the system, for instance by increasing protein concentration in the reactor, omitting glycerol^4^, or further optimizing the exact composition of the feeding solution^6^, will render even higher protein production rates possible.

Although this specific microfluidic device is designed to study systems requiring non-equilibrium conditions such as synthetic cells and biomolecular oscillators that operate out-of-equilibrium, the methods described here could also be expanded to be used for applications that would benefit from a higher protein synthesis yield, including for instance continuous dialysis protein synthesis combined with on-chip purification^30,31^ or protein crystallization^12^. Besides, the simplicity and time-efficiency of the silanization protocol and method to prepare PEG-DA hydrogel membranes by utilizing pneumatic valves presented here also suggests itself for the implementation of other applications that involve hydrogels^31,32^ in-double-layer microfluidic devices, such as on-chip concentration gradient generator- and separation devices^31^.

## Methods

### Microfluidic chip fabrication

The device with 8 reactors and 9 fluid inputs (Figure 1, and Supplementary Figure 1) is based on a previous design^2,17^. Molds for the control, the flow layer, and the anti-evaporation layer were fabricated on separate wafers by standard photolithography techniques. The positive photoresist AZ 10XT-60 (Merck) with a height of 14 μm was used to generate the flow channel features. For the control layer and the anti-evaporation layer, SU-8 photoresist (Microchem 3025, Kayaku Advanced Materials) was used to generate the channel features with a height of 40 μm. Subsequently, each of the wafers was treated with trimethylchlorosilane. For the flow and the anti-evaporation layers, PDMS with an elastomer to crosslinker ratio of 5:1 was prepared and poured over the wafers. The wafers coated with PDMS were placed in a desiccator for 40 min prior to baking. For the control layer, PDMS with a 20:1 elastomer to crosslinker ratio was spin-coated at 1400 rpm onto the wafer and left to relax for 40 min prior to baking. The flow and control layers were partially cured at 80°C for 20 minutes. The flow layer was then cut out and aligned onto the control layer. The aligned devices were placed back in the oven at 80°C for 90 minutes. The anti-evaporation layer was cured at 80°C for 90 minutes. After the curing, the devices and the anti-evaporation layers were removed from the wafer and the inlets were punched. The aligned devices were plasma bonded to a glass slide and subsequently the evaporation layer was plasma bonded on top of the chip.

### Chip silanization and formation of PEG-DA hydrogel membrane

To prime the chip, control lines were filled with water and pressurized with air at 1.38 bar, except for the hydrogel-forming valves, which were pressurized with nitrogen at 1.38 bar. The inner walls of the flow channels, except for the reactors, were oxidized by flowing freshly prepared solution of H_2_O:H_2_O_2_ (30% (v/v)):HCl (36%) solution in a ratio of 5:1:1 v/v for 10 minutes, followed by a washing step with MilliQ water for 4 min. The silanization solution was freshly prepared by mixing MilliQ water with 0.4% (v/v) 3-(Trimethoxysilyl)propylmethacrylat (TMSPMA, Sigma) and 0.4% (v/v) glacial acetic acid (VWR) and let flow for 20 min through the oxidized channels. Subsequently, the channels were washed with MilliQ water for 15 min.

Meanwhile, an aqueous solution for the polymerization of the hydrogel membrane was prepared. First, the PEG-DA (M_n_ 700 g/mol, Sigma) was mixed with photo-initiator 2-hydroxy-2-methylpropio-phenone (Sigma) (90/10% w/w). Subsequently, different volumes of UltraPure water (Invitrogen) were added to the mixture, as specified in section **Characterization of PEG-DA hydrogel membranes**. The solution was perfused with nitrogen gas for up to 20 min to remove oxygen. However, we found that shorter times of less than 1 min are sufficient for the hydrogel formation. After the silanization process, the anti-evaporation layer was pressurized with nitrogen at 1.38 bar to deplete oxygen from the chip. The PEG-DA solution was injected into the feeding chamber for 10 min, after which the hydrogel-forming valves were closed, and the remaining PEG-DA was removed by washing with water. The remaining PEG-DA was polymerized for 2 × 30 s using an Omnicure S1500 200W UV curing lamp with a standard filter (320 nm – 500 nm). After the formation of the membranes, the hydrogel forming valves and the anti-evaporation layer were disconnected from the nitrogen source. The anti-evaporation layer was primed with water and the formed membranes were washed with UltraPure water for 20 hours and subsequently passivated with bovine serum albumin (1%, Sigma-Aldrich) in (50 mM HEPES, 100 mM potassium chloride, 10 mM magnesium chloride) for 1-3 hours, and finally primed with a wash buffer (50 mM HEPES, 100 mM potassium glutamate, 11.8 mM magnesium acetate).

### Solute diffusion through the hydrogel membranes

The reactors were thoroughly washed and filled with the wash buffer, while the feeding channel was loaded with either 10 μg/mL of methylene blue, Fluoresceinisothiocyanat-dextran (FITC-dextran, 10 kDa) or Tetramethylrhodamine-dextran (TMR-dextran, 40 kDa) solutes. The peristaltic pump was actuated at 20 Hz to mix the solutions inside the reactor. The reactor was imaged every 10 min.

### Energy and feeding solution preparation

The energy solutions were prepared as described previously^29^ although at a higher final concentration (4x). The 4x energy solution contained 1.2 mM of each amino acid, 47.2 mM magnesium acetate, 400 mM potassium glutamate, 8 mM ATP and GTP, 4 mM CTP, UTP and TCEP (tris(2-carboxyethyl)phosphine hydrochloride), 14 mg/mL tRNA, 80 mM creatine phosphate, 0.08 mM folinic acid, 8 mM spermidine, and 200 mM HEPES. The prepared energy solution was then diluted to 2.5x working concentration, or used in a feeding solution. The feeding solutions were prepared by combining the energy solution or Solution A (PURExpress) at the desired concentrations with PURE buffer (final concentration in the feeding solution: 13 mM HEPES, 1.4 mM magnesium acetate, 1.7 mM magnesium chloride, 7 mM potassium glutamate, 4.2 mM potassium chloride, 4 mM TCEP, 2.8 mM 2-mercaptoethanol).

### Device setup for cell-free expression

PURExpress solution B supplemented with RNase inhibitor (2 U/μL), mScarlet, and TCEP (10 mM), PURExpress solution A or energy solution 2.5x, and DNA solutions were mixed in the microfluidic reactors on the microfluidic chip in a 2:2:1 v/v ratio. The DNA solution at five times its final concentration was prepared by mixing 10 nM eGFP linear templates and 6.25 μM Chi DNA as described previously^2^.

For continuous dialysis batch experiments, the reactor was imaged every 10 minutes and the feeding channel solution was replenished if indicated. For chemostat experiments, the reactor was imaged every 15 min, a 20% fraction of the reactor volume was replaced with fresh components with the same 2:2:1 ratio every 15, 30, or 60 min, while the feeding channel solution was replenished every 15 min, if indicated. Details on the operation of the microfluidic chip can be found in Supplementary Tables 1 and 2.

### Data acquisition and analysis

Solenoid valves, microscope, and camera were controlled by a custom Matlab and LabVIEW program. The chip and microscope stage were enclosed in an environmental chamber at 34 °C. The fluorescence was monitored over time on an automated inverted fluorescence microscope (Nikon), using 20x magnification and FITC / Cy5 / mCherry filters. The microscope hardware details are as described previously^17^. Fluorescence images were analyzed using a custom Python script.

## Acknowledgements

The authors would like to express their gratitude to Nadanai Laohakunakorn and Gregoire Michielin for helpful input and comments. This work was supported by a Human Frontier Science Program Grant RGP0032/2015; the European Research Council under the European Union’s Horizon 2020 research and innovation program Grant 723106; and a Swiss National Science Foundation grant (182019).

## Author Contributions

B.L. & L.G. performed experiments. B.L., L.G. and S.J.M. designed experiments, analyzed data, and wrote the manuscript.

## Notes

The authors declare no competing interests.

## Supplementary information

**Supplementary Figure 1.**
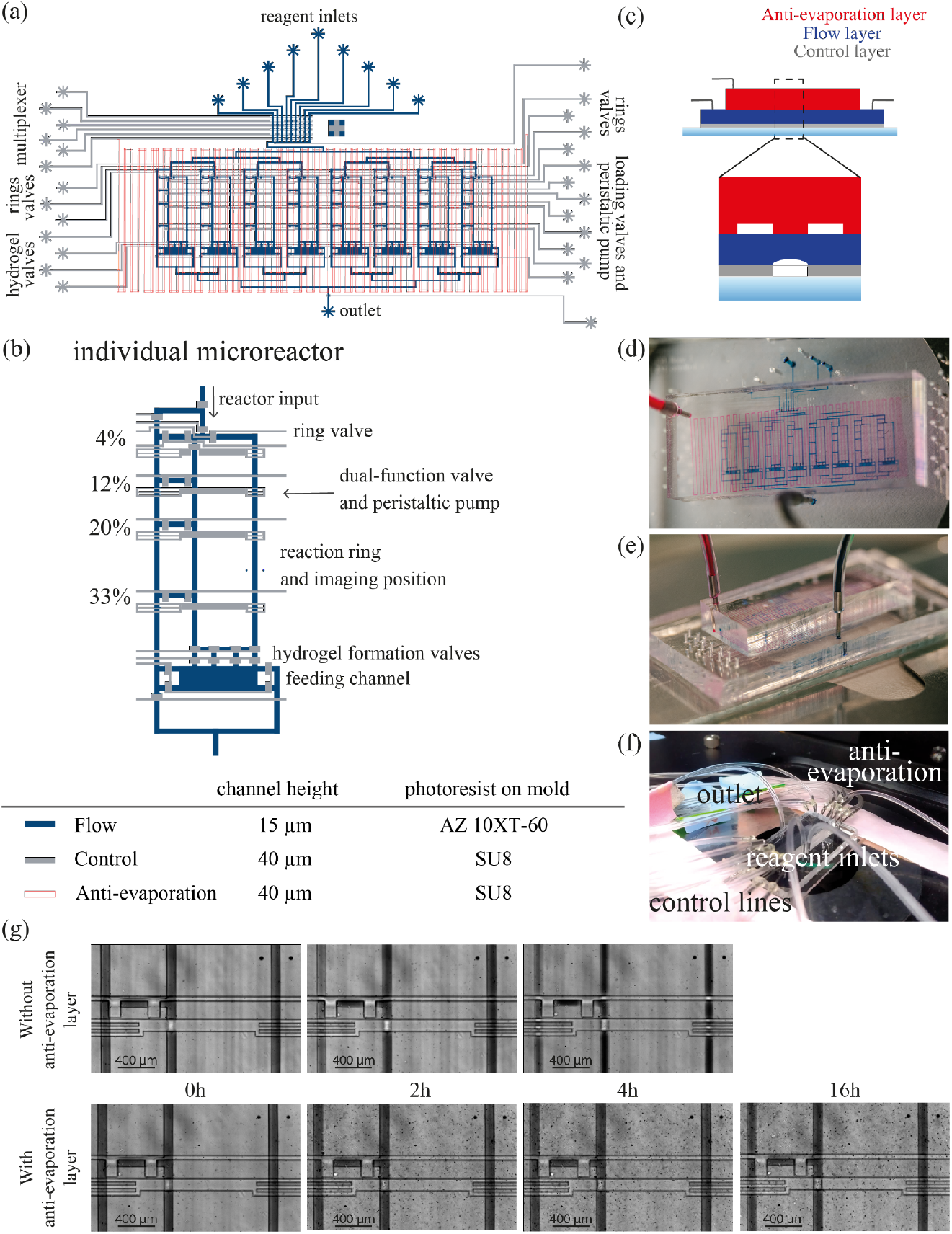
Design of the microfluidic device with hydrogel membranes:(a) Design schematic of the microfluidic device. The control layer is shown in gray, the flow layer in blue, and the anti-evaporation layer in red. The device features eight chemostat reactors. (b) Close-up of a microfluidic reactor and table of channel heights and corresponding photoresists used during mold fabrication. Each reactor has four outlets corresponding to four different dilution fractions. Four Control lines serve dual-functions as valves and peristaltic pumps. The width of a flow channel is 100 μm. (c) Schematic depiction (not to scale) of the microfluidic chip cross-section. Image of the microfluidic chip from the top (d) and from the side (e), for visualization the flow channels and the anti-evaporation channel are filled with blue and red dye, respectively. (f) Image of microfluidic chip connected to the control lines, reagent inputs and outlet. (g) Comparison of the water evaporation in the chip with and without the anti-evaporation layer at different times at 34°C. The flow channels were filled with blue dye for better visualization.

**Supplementary Figure 2.**
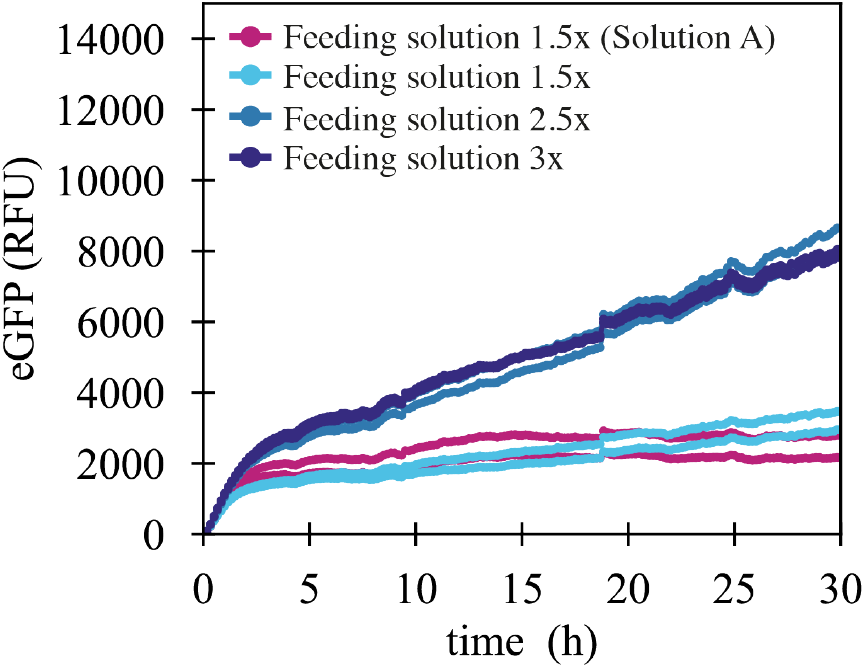
Time course of in vitro eGFP expression levels in batch with continuous dialysis reactions for different feeding solution concentrations based on the commercial solution A (PURExpress) or home-made energy solution. Each curve represents a technical replicate.

**Supplementary Figure 3.**
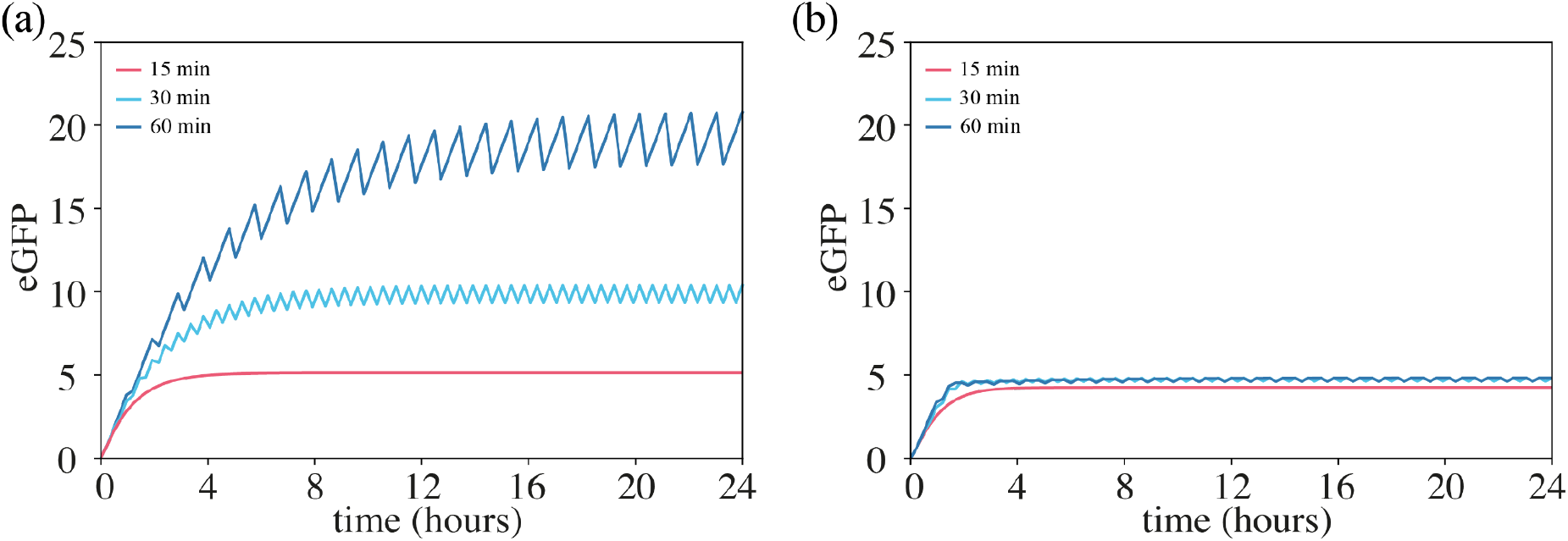
Chemostat simulations: The chemostat was simulated using a minimal resource-dependent TX-TL model^2^ periodically diluting and replenishing species and solving ODEs every 15 min to match the experimental measurements. The resource R_0_ was chosen so: (a) the resource depletion is not rate limiting (R_0_ = 1000), (b) the resource depletion is rate limiting (R_0_ = 30). 20% of the chemostat reactor volume was diluted at different time intervals of 15, 30, and 60 min. Parameter conditions α = 0.7, β = 0.07, K = 1 and initial conditions pT = 10, dG = 2, with all other species set to zero.

**Table 1.** Microfluidic chip operations for batch, batch with static dialysis and batch with continuous dialysis reactions

| Batch reaction without dialysis |  |  |
| --- | --- | --- |
| Step | Operation | Solution |
| **0A Initial fill** |  |  |
|  | Flush rings | PURE solution |
|  | Load 20% (Flush through outlet 20%) | eGFP DNA |
| **0B** | **Mix** |  |

| Batch reaction with static dialysis |  |  |
| --- | --- | --- |
| Step | Operation | Solution |
| **0A Initial fill** |  |  |
|  | Flush rings | PURE solution |
|  | Load 20% (Flush through outlet 20%) | eGFP DNA |
| **0B Feeding chamber fill** |  |  |
|  | Load feeding chamber | Feeding solution |
| **0C** | **Mix** |  |

| Batch reaction with continuous dialysis |  |  |
| --- | --- | --- |
| Step | Operation | Solution |
| **0A Initial fill** |  |  |
|  | Flush rings | PURE solution |
|  | Load 20% (Flush through outlet 20%) | eGFP DNA |
| **0B Feeding chamber fill** |  |  |
|  | Load feeding chamber | Feeding solution |
| **0C** | **Mix** |  |
| Repeat the following steps every 10 min |  |  |
| **1A Feeding chamber fill** |  |  |
|  | Load feeding chamber | Feeding solution |

**Table 2.**
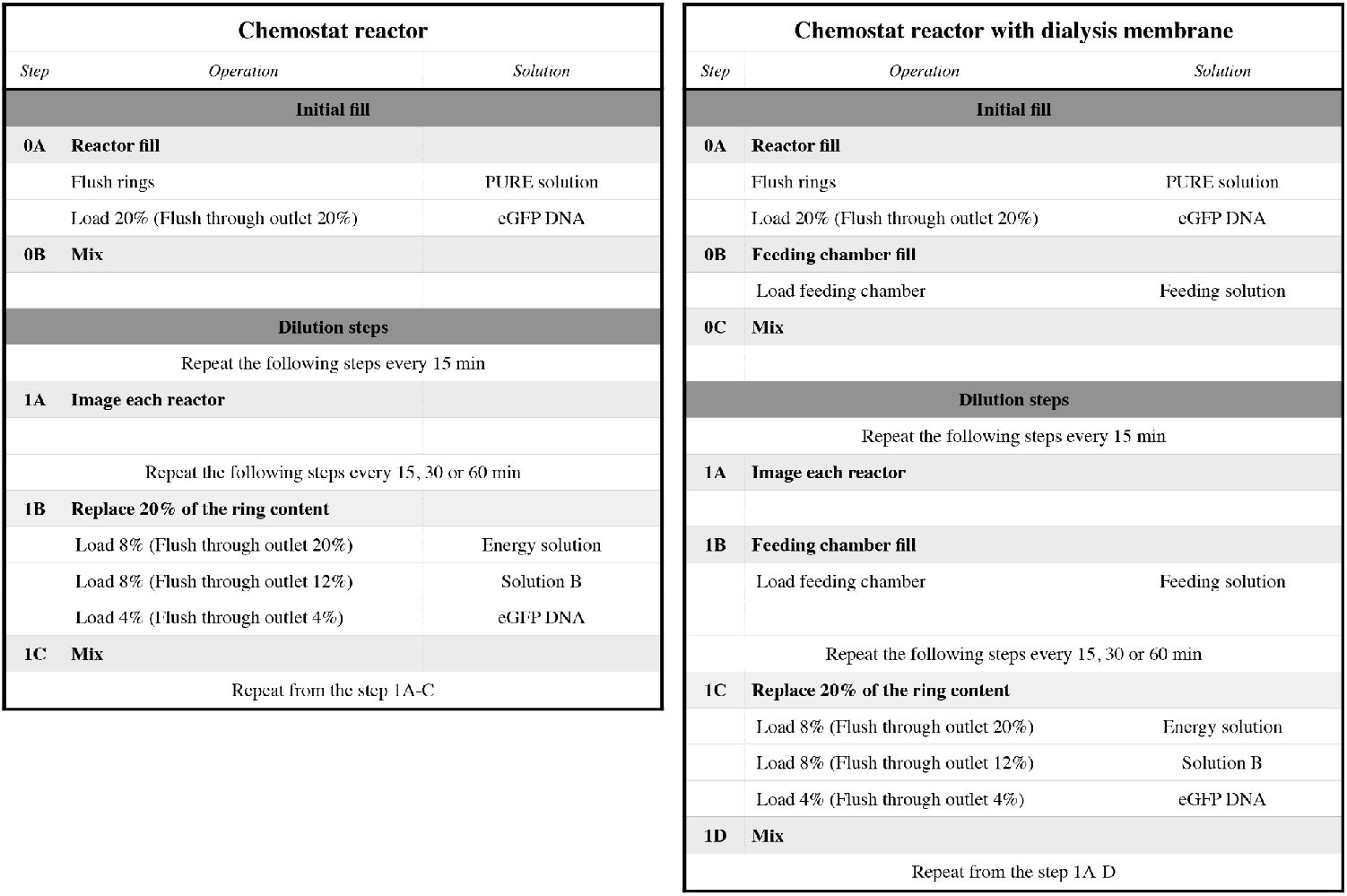
Microfluidic chip operations for steady-state reactions

## Notes

### Competing Interest Statement

The authors have declared no competing interest.

